# Establishing wonder oil, Solanesol, as a novel inhibitor for Focal Adhesive Kinase by in silico strategies

**DOI:** 10.1101/086660

**Authors:** Betty Daneial, Devashish Das, V. J. Jacob Paul, R. Guruprasad

## Abstract

Focal adhesion kinase (FAK) plays a primary role in regulating the activity of many signaling molecules. Increased FAK expression has been implicated in a series of cellular processes, including cell migration and survival. Inhibiting the activity of FAK for cancer therapy is currently under investigation. Hence, FAK and its inhibitors has been the subject of intensive research. To understand the structural factors affecting inhibitory potency, kinetic analysis, molecular docking and molecular dynamics simulation were studied in this project. Though, Solanesol was found have inhibitory activities towards FAK, no *in silico* tests were ever done on the same.

Due to high flexibility of Solanesol (Rotatable bonds = 25), it is difficult to analyze using normal docking protocols. This paper introduces a novel method to dock and analyze molecules with high flexibility based on weighed contact based scoring method. This method uses blind docking technique, which was developed for protein peptide docking method, to generate conformations which were used to calculate contact based weights of residues. This method reveals the possible binding site for the small molecule. An exhaustive docking search on the acquired area reveals the docked confirmation of the compound. The final docked conformation was subjected to molecular dynamics to understand of binding stability. This study is in a good agreement with experimental results which shows Solanesol binds at ATP binding site and inhibit the phosphorylation of Focal Adhesion Kinase.

## Background

### Focal Adhesion Kinase

Among the 10 hallmarks of cancer, “Tissue Invasion and Metastasis”, is the most devastating one, as it allows the oncogenic cells to migrate to new areas in the body with more open space and more nutrients. It significantly reduces survival rates and prognosis for patients. The loss of cellular adhesion to the extracellular matrix regulation can lead to increased cellular proliferation, decreased cell death, and altered cellular differentiation status and cellular migratory capacity (Hanahan & Weinberg, 2000; Hanahan & Weinberg, 2011; Lazebnik, 2010; van Nimwegen & van de Water, 2007).

The FAK4 family kinases, including Focal Adhesion Kinase (FAK), regulate cell adhesion, migration, and proliferation in a variety of cell types. Focal adhesions are heterodimeric-transmembrane integrin receptors located within sites of close opposition to the underlying matrix which mediate adhesion of cells to extracellular matrix. TYR397 position FAK phosphorylation simulated by Integrin engagement and clustering creates high affinity binding site for SRC and SRC family kinases. In turn, FAK-SRC complex phosphorylates many components of focal adhesion, which result in changes in initiation of signaling cascades and adhesion dynamics. Other than the FAK catalytic activity, FAK additionally functions as a scaffold to organize signaling and structural proteins within focal adhesions (Cohen & Guan, 2005; Guan, 1997).

Alteration in FAK expression not only have been associated with tumorigenesis and increased metastatic potential, it is also reported to cause multiple cancers, including colon, breast, thyroid, prostate, cervical, ovarian, head and neck, oral, liver, stomach, sarcoma, glioblastoma, and melanoma (Guan, 1997).

### Solanesol

Solanesol is a polyisoprenoid alcohols or polyprenols, found mainly accumulates in solanaceous crops, including tobacco, tomato, potato, eggplant, and pepper plants. In industries, Solanesol is extracted commercially from it richest source, tobacco plant.Commercially, Solanesol is widely used as an intermediate for the synthesis of ubiquinone drugs, such as coenzyme Q10 and vitamin K2 as well as Vitamin K and Vitamin E. It known to possess activities like antibacterial, antifungal, antiviral, anticancer, anti-inflammatory, and anti-ulcer activities, and its derivatives also have anti-oxidant and anti-tumor activities, in additionto other bioactivities (Yan et al., 2015; Srivastava et al, 2009; Suzuki, Tomida & Nishimura, 1990; Tomida & Suzuki, 1990; Zhao, Zu, Li, & Tian, 2007; Severson, Ellington, Schlotzhauer, Arrendale, & Schepartz, 1977). Derivatives of Solanesol, (S)-2,3-dihydropolyprenyl, monophosphate, and agents for inhibiting the metastasis of cancers (Okamoto, Tsuji, & Yamazaki, 1994).

### Objective

The main aim of this paper is to establish Solanesol as a Focal adhesion kinase inhibitor by the means of *in silico* methods. Though Solanesol used as bioactive agent in industries for decades, due to its highly flexible nature, there is no successful *in silico* protein binding and simulation data available online till date.

Solanesol has 25 rotatable bonds which makes very difficult to dock directly to the protein structure by conventional method. Although there are various methods for prediction of binding site in FAK, the flexible nature of Solanesol as well as the size of the compound makes it difficult to fit the active site. For the purpose of predicting the actual binding site, blind docking method can be used. Blind docking method was introduced for the purpose of docking peptide molecules to the protein molecule but is a tested method for binding of small molecules when binding site is unknown (Hetényi & van der Spoel, 2006).

Method of blind docking comprise of locking of ligand molecule’s all the torsions, for the purpose of reducing calculation time, and then docking on the complete protein surface. Then the best cavity or binding pocket is selected, based on the clustering of highest binding affinity conformations (Hetényi & van der Spoel, 2006; Hetényi & van der Spoel, 2002).

In this paper we propose a method of binding of highly flexible compound to protein targets with using an enhanced contact based scoring method. This method scores the residues rather than the conformations. The higher scored residues where then used for more “focused” docking on those residue region.

## Materials and methods

### Protein selection and preparation

The crystallographic co-ordinates for Focal adhesion kinase (PDB ID: 4Q9S) (George et al, 2014) were retrieved from the Protein Data Bank (PDB). Prior to docking, protein structures were prepared by removing water molecules using UCSF Chimera software (Pettersen et al, 2004). Following which, bond orders were assigned, and hydrogen atoms were added to the crystal structures.

### Ligand preparation

Solanesol exist in both cis and trans states, for this experiment we considered only trans as it is only found in natural sources (Roe, Oldfield, Geach, & Baxter, 2013). The structure of Solanesol was obtained from PubChem compound (CID 5477212) (Kim et al, 2015; NCBI 2016). Gaussian 09 program (Frisch et al, 2009) was used to obtain the optimum geometry of the structures using the density function theory at the B3LYP/6-31G (d,p) level.

### Molecular docking

All the molecular docking studies of Solanesol to FAK were performed using Autodock 4.2 (Morris et al, 2009). Autodock uses a semi-empirical free energy force field to evaluate binding conformations of ligand while docking. The AutoDockTools (Morris et al, 2009) was used for preparing protein and ligand parameters files.

### Binding site analysis

Solanesol is a 45 carbon chain with 26 rotatable bonds. As it is extremely flexible it is hard to determine the bind mode of it with the protein. The commonly used protocol for determination of binding pocket is “Blind docking”, which was initially developed for to determining peptide docking with protein (Hetényi & van der Spoel, 2006; Hetényi & van der Spoel, 2002). In this method the constrained ligand (or peptide) is docked with the whole protein surface. The place where it forms a cluster with higher energy determines the binding site. Then these sites were used for “refined docking” where the lowest binding modes for each of these places (in case if there are more than one) where determined by molecular mechanics and molecular dynamics studies.

For calculating the possible area of interaction or binding site of a highly flexible ligand we enhanced blind docking using Ligand Contact Based Scoring function for Residues (LCBSR). This is an atomic contact\clash based scoring method, in which the residues are scored based on the higher favorable interactions and probability of formation of a hydrogen bond. As it doesn’t depend on the clustering or solely on binding energy, it statistically enhances the probability of finding the possible binding site.

It can be represented as,

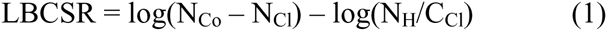

Where, ligand contact based scoring of residue (LCBSR) is calculated using “Number of Contacts” (NCo) which is the number of occurrences where atoms of residue (r) are in contact with any atom of the ligand in all the conformations. “Number of Clashes” (NCl) is the number of occurrences when atoms of residue (r) are in clashes or unfavorably overlapped with any atom of the ligand in all the conformations. “Number of Hydrogen Bonds” (NH) is the number of occurrences when atoms of residue (r) are forming a hydrogen bond with any atom of the ligand in all the conformations. “Number of Clashes” (CCl) is the number of conformers where residue (r) is in an unfavorable overlap with any atom of the ligand.

The “Contact” here is defined as the instance when the difference between the distance of two atoms and the sum of their van der Waal radii is 0.4 A or more or, in other words, the distance is greater than the sum of van der Waal radii (Eq 1) of two atoms. Whereas “Clash” is defined as the condition where the van der Waal radii of two atoms unfavourably overlaps each other and the distance is lower than the sum of radii (Eq 2) of the two atoms. This can be represented as:

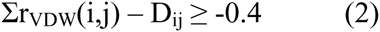

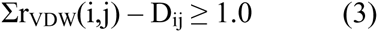

Where, *∑r_VDW_* (*i*, *j*) is the sum of van der Waal radii of interacting atoms of ligand (i) and residue (j) and D_ij_ is the distance between interacting atoms of ligand (i) and residue (j). A Higher value of LCBSR of any residue implies the residue may be a part of the binding pocket for the ligand and, if it is capable, it may also form a hydrogen bond with the ligand. Lesser score implies a lower interaction or high chances of unfavorable clashes or low chances of forming a hydrogen bond.

## Experimental design

### Blind docking

For this experiment, Solanesol was docked a total of 3 times (Table 1) with “Blind Docking” protocol. For getting an unbiased result, all the rotatable bonds were kept unconstrained for all the experiments. The protein was covered using a 126 × 126 × 126 grid box with protein centre as grid centre. For experiment ligand starting position was changed.

**Table 1:** Blind docking analysis with three different experiments (with different starting conformations) using standard Autodock protocol

| Experiment | Total Conformations | Grid Center | L.B.E ($\Delta G$) in kCal/Mol | Total Cluster | Highest number of member in any cluster | Average B.E of cluster with most number of members |
| --- | --- | --- | --- | --- | --- | --- |
| Blind Docking I | 500 | Center of Protein | -7.08 | 328 | 10 | 0.23 |
| Blind Docking II | 200 | Center of Protein | -8.54 | 160 | 5 | -0.19 |
| Blind Docking III | 200 | Center of Protein | -5.68 | 187 | 3 | 0.13 |

The docking resultant file from Autodock was then converted into multiple PDB files using Autodock scripts. All the contacts and clashes, as well as hydrogen bonds between the conformation and protein, were calculated using UCSF Chimera tool.

Considering all the conformers may lead to false positives, thus conformers were separated in three criteria:

1. Binding energy less than −2.0 kcal/mol
2. Binding energy less than −3.0 kcal/mol
3. Binding energy less than −4.0 kcal/mol.

Provided −4.0 is roughly half the average of the binding energy (Average B.E = −7.10 kCal/mol) of all the three experiments, ensuring only conformations with low B.E were considered. The resultant values from each of these three were added to corresponding residues. This way the residues interacting more with the conformations of better binding energy will have a better score.

For getting a statistically significant result, all the scores of the residues from all the three experiments were added to get the final score of each residue. Only those residues which appeared in more than 2 experiments were considered.

### Refined docking

A binding site was formed using residues with a higher LCBSR score (percentile = 0.50). This binding site was then used for refined docking using Autodock. The experiment was done twice with (1) relaxed parameters, GA maximum energy evaluations 2.5 × 106, for 200 GA runs (2) exhaustive parameters, GA maximum energy evaluations 3.5 × 107, for 200 GA runs.

### Knowledge-based rescoring

All conformations were rescored using DSX, Drug Score eXtended, (Neudert & Klebe, 2011), knowledge-based rescoring based on the DrugScore formalism, to estimate the affinity of conformation for FAK. The best conformations were selected based on the rescored values. The best conformation bound complex of FAK was further used for molecular dynamics simulation studies.

### Molecular dynamics

Molecular dynamics simulations for FAK protein as well as Solanesol bound FAK were performed using the GROMACS (Groningen Machine for Chemical Simulations) 4.6 (Hess, Kutzner, Van Der Spoel, & Lindahl, 2008) software with GROMOS96 (53a6) force field. PRODRG (Van Aalten et al, 1996) server was used to generate topology files for Solanesol. Charges were kept full and no energy minimization was done using PRODRG.

The complex was solvated in a dodecahedron box with SPC model water model molecules and periodic boundary conditions were used. One negatively charged chlorine ion (Cl-) was added to the system for maintaining the system’s neutrality.

The Lincs and Shake algorithm (Hess, Bekker, Berendsen, & Fraaije, 1997) were used for constraining bond length and fixing all bonds containing hydrogen atoms respectively.

For electrostatic calculations, Particle Mesh Ewald (PME) (Darden, York, & Pedersen, 1993) method was used, with a coulomb cutoff of 1.2nm, Fourier spacing of 0.16 nm and an interpolation of order 4. Energy minimization of the system was carried out using steepest descent algorithm with a tolerance value of 1000 kJ mol-1nm-1. After energy minimization, NVT and NPT equilibrations were done on the system until it reached the room temperature and water density. Production MD was performed for 20 ns time duration for both the simulations.

### Molecular dynamics trajectories analysis

Root mean square deviation (RMSD) and root mean square fluctuations (RMSF) of FAK backbone were calculated using “g_rms” and “g_rmsf” utility commands, respectively. A spherical probe of radius 1.4 Å across the protein surface was used for calculating solvent-accessible surface area (SASA) by “g_sasa” tool of Gromacs. Hydrogen bonds between Solanesol and FAK were calculated using “g_hbond” tool with proton donor and acceptor distance ≤ 3.5 Å and the angle between acceptor-donor-hydrogen ≤ 30.0 degrees.

### Binding free energy calculations

Molecular mechanics Poisson-Boltzmann surface area (MMPBSA) (Massova & Kollman, 2000) approach was to estimate the binding free energy of protein-ligand interaction. For this purpose, “g_mmpbsa” (Kumari, Kumar, & Lynn, 2014) tool was used. The tool calculates the molecular mechanics potential energy and the free energy of solvation and excludes the entropy calculations. MM-PBSA calculations were performed using 1000 snapshots taken from last 5 ns of trajectories of the complex system.

## Result & Conclusion

### Docking Analysis

#### Blinding docking and LBCSR Score

Solanesol was docked against FAK structure (PDB id: 4Q9S) using Autodock4.2 three times (as mentioned in materials & methods). For all the three times, a grid of size 126 × 126 × 126 with 0.375A [Figure 1] spacing was created with protein center (Centre coordinates = 9.7, 0.16, 15.1) as the grid center, as per the normal “blind docking” protocol. Dielectric constant value was kept default.

**Figure 1:**
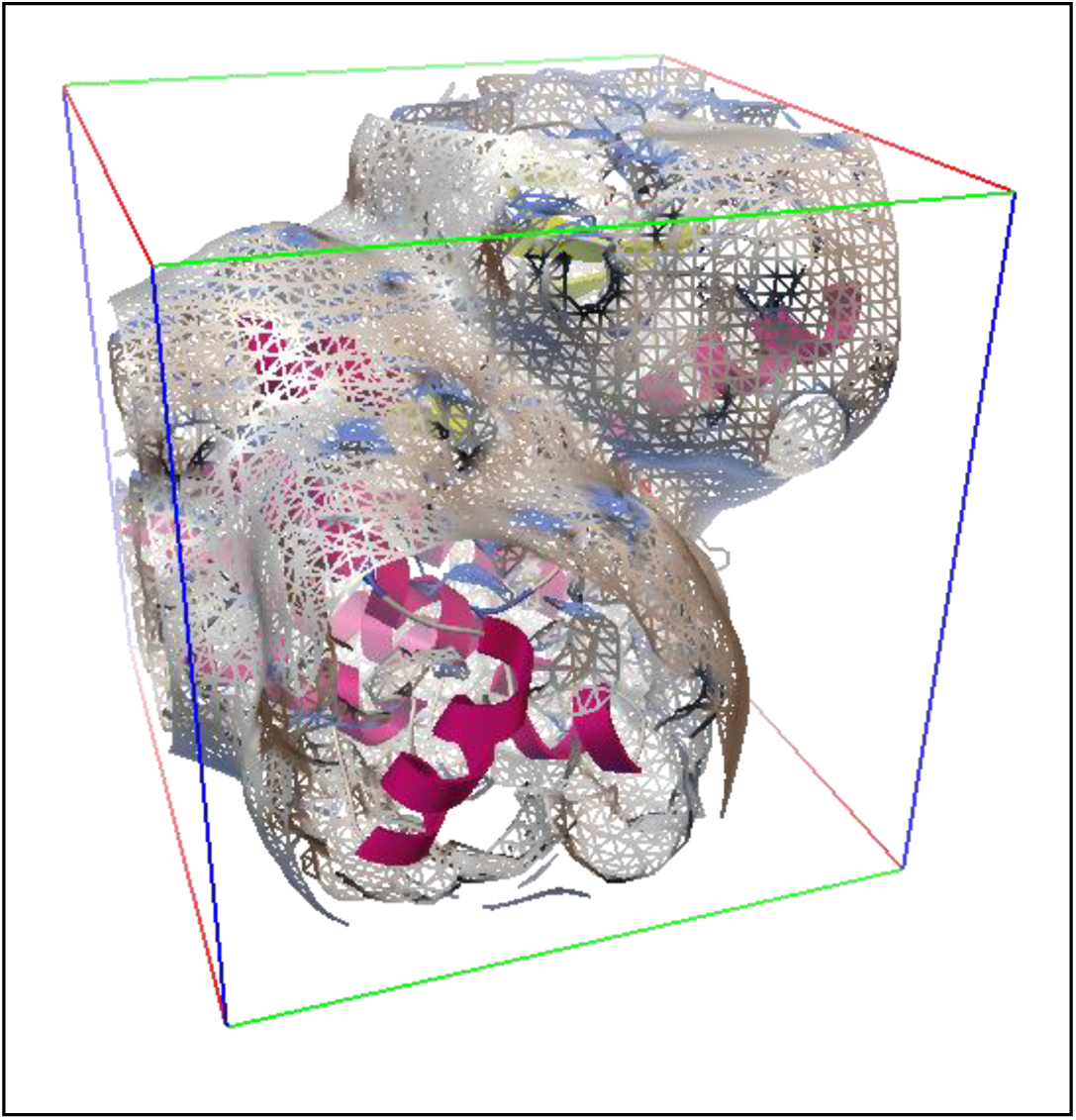
Autogrid map on Focal adhesion kinase as generated by Autogrid4 when blind docking is done on the whole surface.

Solanesol shows very small cluster with insignificant binding affinity towards FAK when docked “blindly” [Table 1]. Though, the structure show quite high binding affinity towards the kinase but the conformations with higher binding energy fails to form any significant cluster.

From all the generated conformations of Solanesol which were having binding energy lower than-4.0 kCal/mol, all the favorable and unfavorable overlaps of the atoms were calculated using “FindClash” tool of UCSF Chimera. Chimera tool, “FindHbond” was also used to find the hydrogen bonds between the ligand and residue atoms. The default values were kept for all the calculation in Chimera.

An in-house python script was used to calculate the number of “Clashes”, “Contacts” and “Hbonds” between all the conformations and residues as well as individual scores. The scores from all the three experiments were added to give a final score for each residue [Table 2].

**Table 2:** Top residues with best LBCSR scores of all three experiments

| Residues | Blind Docking I | Blind Docking II | Blind Docking III |
| --- | --- | --- | --- |
| LYS-454 | 5.66 | 16.50 | 11.27 |
| ASP-564 | 8.89 | 22.35 | 14.77 |
| ARG-550 | 7.48 | 16.33 | 13.64 |
| GLU-500 | 2.48 | 12.06 | 6.14 |
| ARG-426 | 3.04 | 12.11 | 3.18 |
| GLU-430 | 5.02 | 13.87 | 13.49 |
| LEU-501 | 4.25 | 11.98 | 8.23 |
| GLY-563 | 4.25 | 12.08 | 6.76 |
| GLU-471 | 6.78 | 22.50 | 9.55 |
| GLN-432 | 5.65 | 21.14 | 11.85 |
| ALA-452 | 4.88 | 12.30 | 7.36 |
| CYS-502 | 5.31 | 12.32 | 9.13 |
| PRO-585 | 4.09 | 15.50 | 7.17 |
| HIS-544 | 4.98 | 15.97 | 9.29 |
| GLY-431 | 5.12 | 17.82 | 11.71 |
| VAL-484 | 3.74 | 8.93 | 5.78 |
| MET-475 | 2.08 | 10.95 | 0.69 |
| ILE-428 | 5.79 | 16.92 | 13.77 |
| MET-499 | 4.42 | 10.30 | 5.78 |
| LEU-584 | 2.30 | 13.20 | 10.42 |
| VAL-436 | 5.55 | 15.14 | 11.36 |
| LEU-553 | 6.30 | 17.18 | 13.47 |

Based on the LBCSR score calculated for all three experiments, the scores for all the residues were plotted as graph (Fig 3) as well as plotted as false colour on the 3D structure of the protein (Fig 2). From the LBCSR score it can be inferred that the ligand shows a high affinity towards the region with Asp564, Asn551, Arg550, Leu553 and Ile428, respectively.

**Figure 2:**
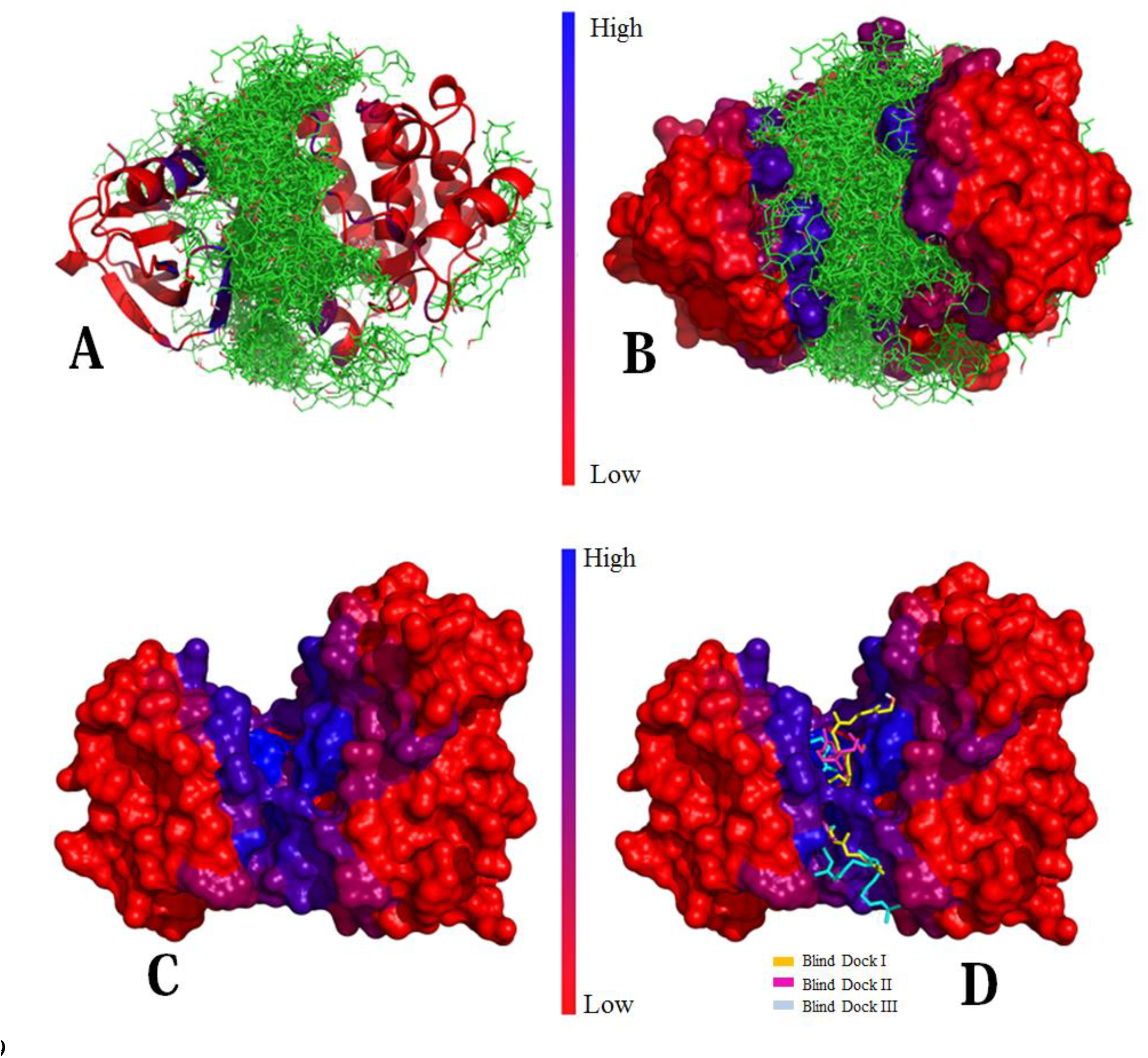
(A,B) FAK structure with colour based on LBCSR score show all the conformations docked “blindly”. (C) Only FAK structure with colour based on LBCSR score (B) Best structures (based on binding energy) from the three docking experiments.

**Figure 3:**
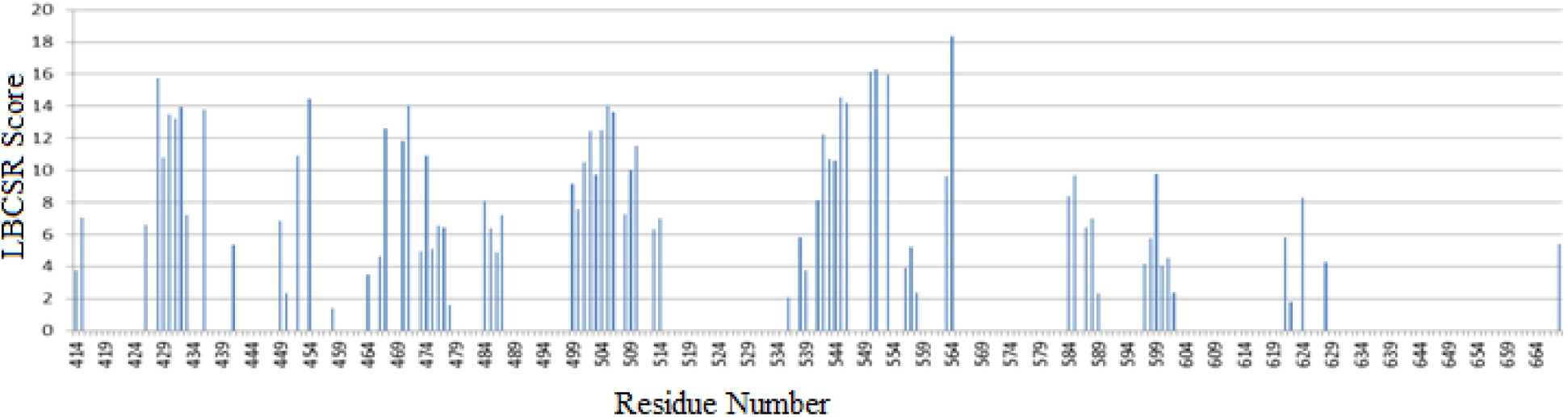
Calculated average LBCSR scores for all the residues of Focal Adhesion Kinase for three experiments.

**Figure 4:**
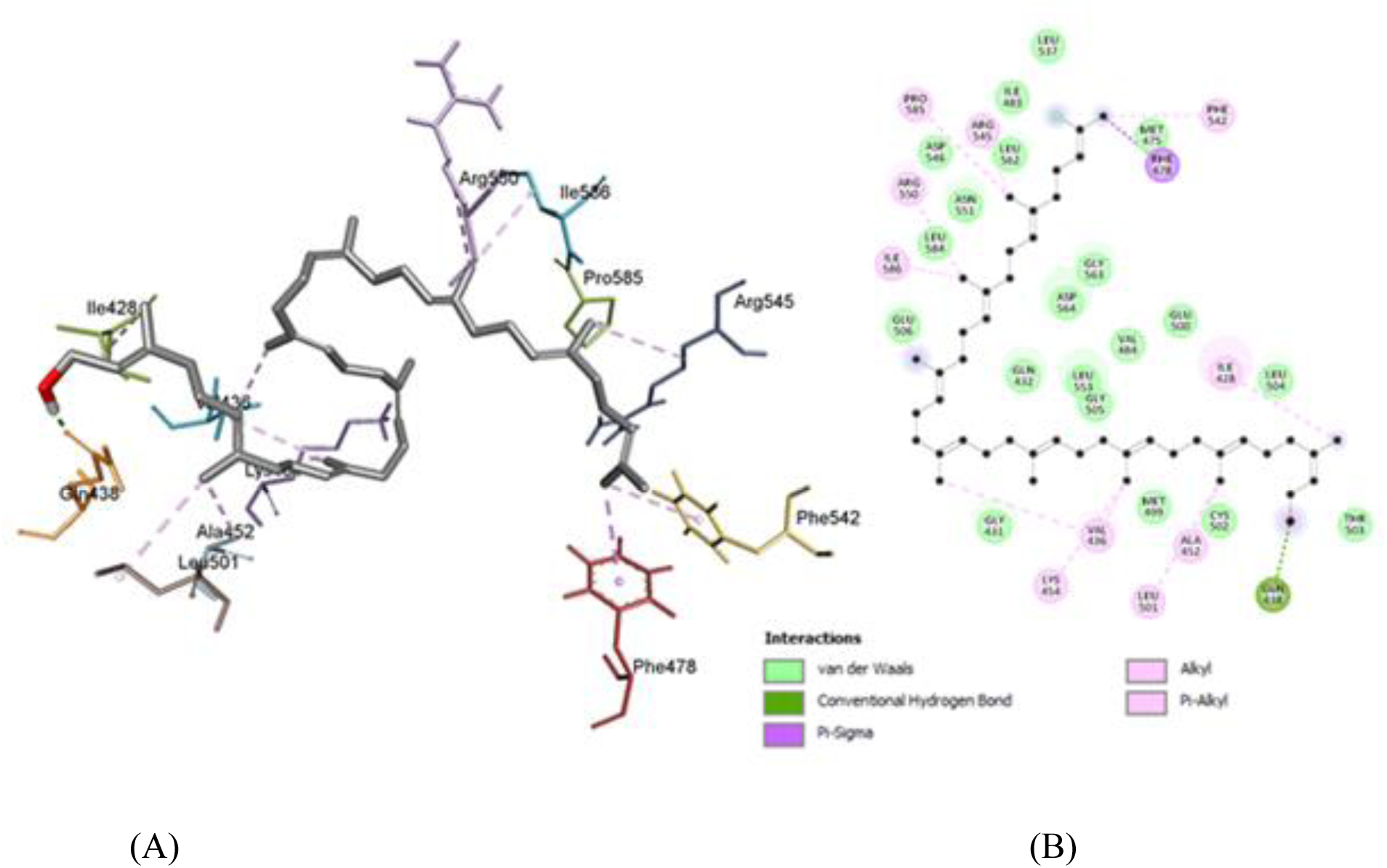
3D (A) and 2D (B) mapof Solanesol when bound to Focal adhesion kinase (FAK)showing different kinds of interactions.

#### Refined docking

All the residues with more than LBCSR score was considered for the active site prediction. A total of 77 residues were found to be above and incidentally which also forms the ATP binding site and the catalytic loop (546-551) and formed between the N and C lobe [1].

Centre of geometry (Coordinates = 10.8, 1.0, 15.0) of these 77 residues were considered for the centre of the binding pocket of Solanesol. A grid of size 60 × 62 × 68 was considered to exactly fit all the 77 residues. Autodock4.2 was again used for docking of Solanesol with FAK with this grid setting for two more times, first time with default setting for 200 conformations and later docked with exhaustive setting for the same number of conformations.

All 400 conformations were rescored using DSX online server with CSD settings. (Table 3)

**Table 3:** DSX, knowledge based scoring of Exhaustive and normal Docking)

|  | Total Score | Per Contact Score | Torsional Score | SAS Score | Binding Free Energy |
| --- | --- | --- | --- | --- | --- |
| Exhaustive Parameter | -166.61 | -0.19 | -11.13 | -16.10 | -7.30 |
| Default Parameter | -131.69 | -0.17 | -12.43 | -8.59 | -7.25 |

### Analysis of Final Docked structure

The hydroxyl end of Solanesol binds to the binding site of ATP and interacts with Ile428, Val436, Ala452, Lys454 and Leu501. These residues which forms the binding pocket for ATP, forms Alkyl-Pi hydrophobic interactions with the double bonds of the ligand. Whereas, the isobutyl end of the ligand, gets attached with the Catalytic loop and αC helix. The ligand interacts with Phe542, Arg545, Arg550 (Catalytic loop) and Phe478 (αC helix terminal) with Alkyl and Sigma-Pi interactions.Pro585, Ile586 near the ATP active site also forms similar hydrophobic interactions with the ligand. Oxygen of Solanesol forms a very conventional H-bond with OE1 of Gln438 which is very near to G-Loop and ATP binding site.[30]

These 12 hydrophobic interactions of 11 residues with Solanesol stabilize the ligand at the middle of N and C lobe of FAK. As the ligand share interacting residues with both the side it may be act as a better inhibitor for FAK. This structure was used for molecular dynamics studies.

### Analysis of Molecular Dynamics

For the measuring the stability, the deviation in the backbone of the protein has been measured relative to time [Fig 5]. After an initial instability, the backbone changes its shapes linearly with a linear slope increase in RMSD between 4.5ns to 7.7ns after again a short destabilization the system vaguely equilibrates from 9.4ns to 14.2ns. It takes the system almost 15ns to equilibrate, after which the system maintains it position and shape. [Fig 5]

**Figure 5:**
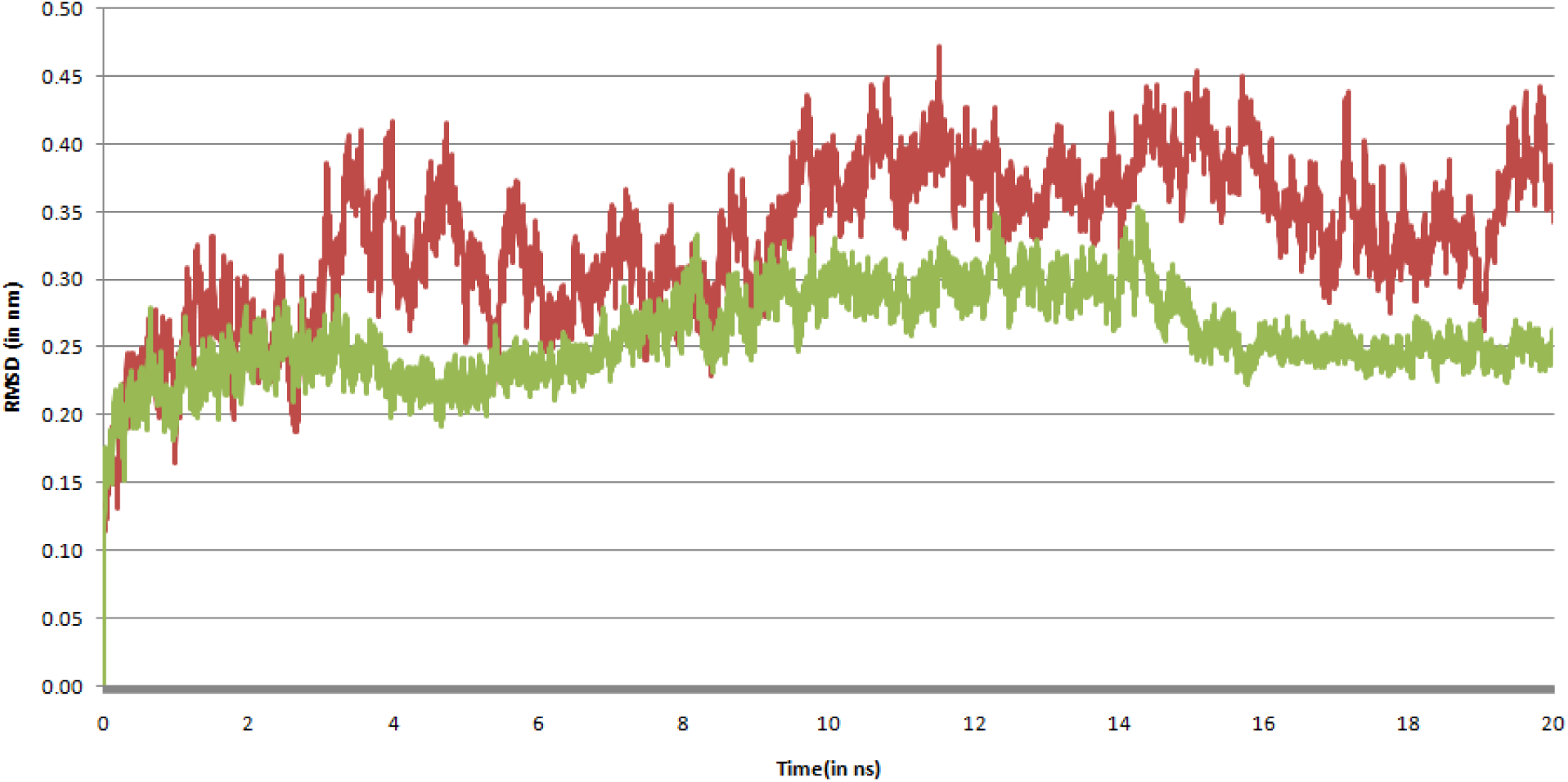
Root mean square deviation (RMSD) of Focal adhesion kinase backbone atoms when bound to Solanesol. It gets stabilized nearly after 16ns of simulation.

Comparison of the RMSD of both the trajectories from the minimized structures shows, Solanesol bound FAK backbone show more stability than that of independent FAK backbone. RMSF (Root mean Square Fluctuation) of Solanesol bond FAK exhibits less fluctuation at the places where Solanesol is bond [Fig 6].

**Figure 6:**
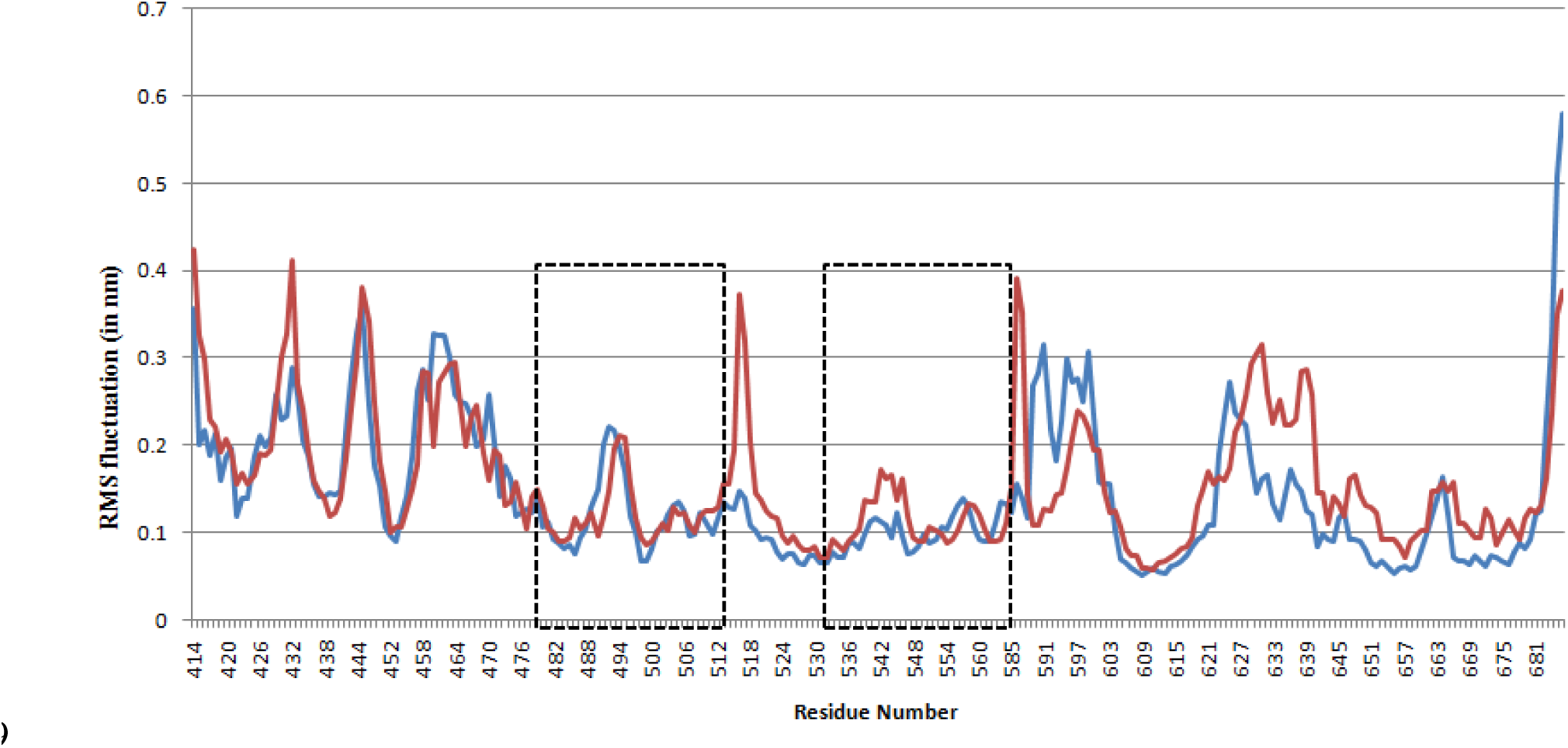
RMS Fluctuation (in nm) in alpha carbon atom of each residue in FAK (Red) and Solanesol bound FAK (Blue). Boxes show the binding site region of FAK.

Total solvent accessible surface area (SASA) was checked by g_sas tool of Gromacs for analyzing the change in surface area with respect to time. It show a quite negative correlation with the RMSD suggesting the better binding of the ligand leads to a lower surface area of the protein thus closing the active site for any further contact. It remains stable for the wider part of 6ns-18ns range after which the backbone gets stabilized and may also affect the surface area.

### MM-PBSA Calculation

Binding site residues for Solanesol were selected by taking 3.5A radius from Solanesol. Molecular mechanics Poisson–Boltzmann surface area (MM-PBSA) was calculated for the last 5 ns with 20 ps steps for the binding site residues versus Solanesol using g_mmpbsa tool.

Per residue analysis of the result was done and plotted using the script provided. This analysis suggests that, Ile428, Gly431, Val436, Val484, Met499, Cys502 and Leu553 interacts most favorably with the ligand. Interestingly these all residues are part of ATP binding pocket of FAK. Gly431 is a part of the G-loop which helps in the phosphorylation of the protein also interacts favorably along with Leu584 which is part of the activation loop.

MM-PBSA based binding affinity (ΔΔG) was calculated using the g_mmpbsa provided script.

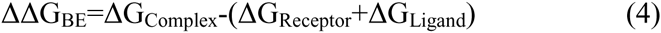

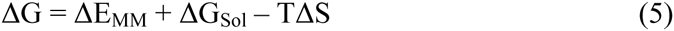

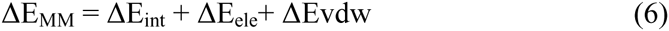

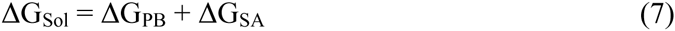

ΔE_MM_, ΔG_Sol_ and TΔS represent the molecular mechanics component in the gas phase, stabilization energy due to salvation, and a vibrational entropy term, respectively. ΔE_MM_ is the summation of ΔE_int_, ΔE_col_, and ΔE_vdw_, which are the internal, coulomb, and van der Waals interaction terms, respectively. ΔG_sol_ is the salvation energy and it is divided into an electrostatic salvation free energy (G_PB_) and a non-polar salvation free energy (G_SA_).

A very low ΔΔG, of −113.85 kJ/Mol, (Table 4) proves Solanesol have a very high binding affinity towards the FAK structure. The change in binding energy with time was also plotted. [Fig 7], It shows that the ligand gets stabilized with a very high binding affinity after 17ns of simulation.

**Figure 7:**
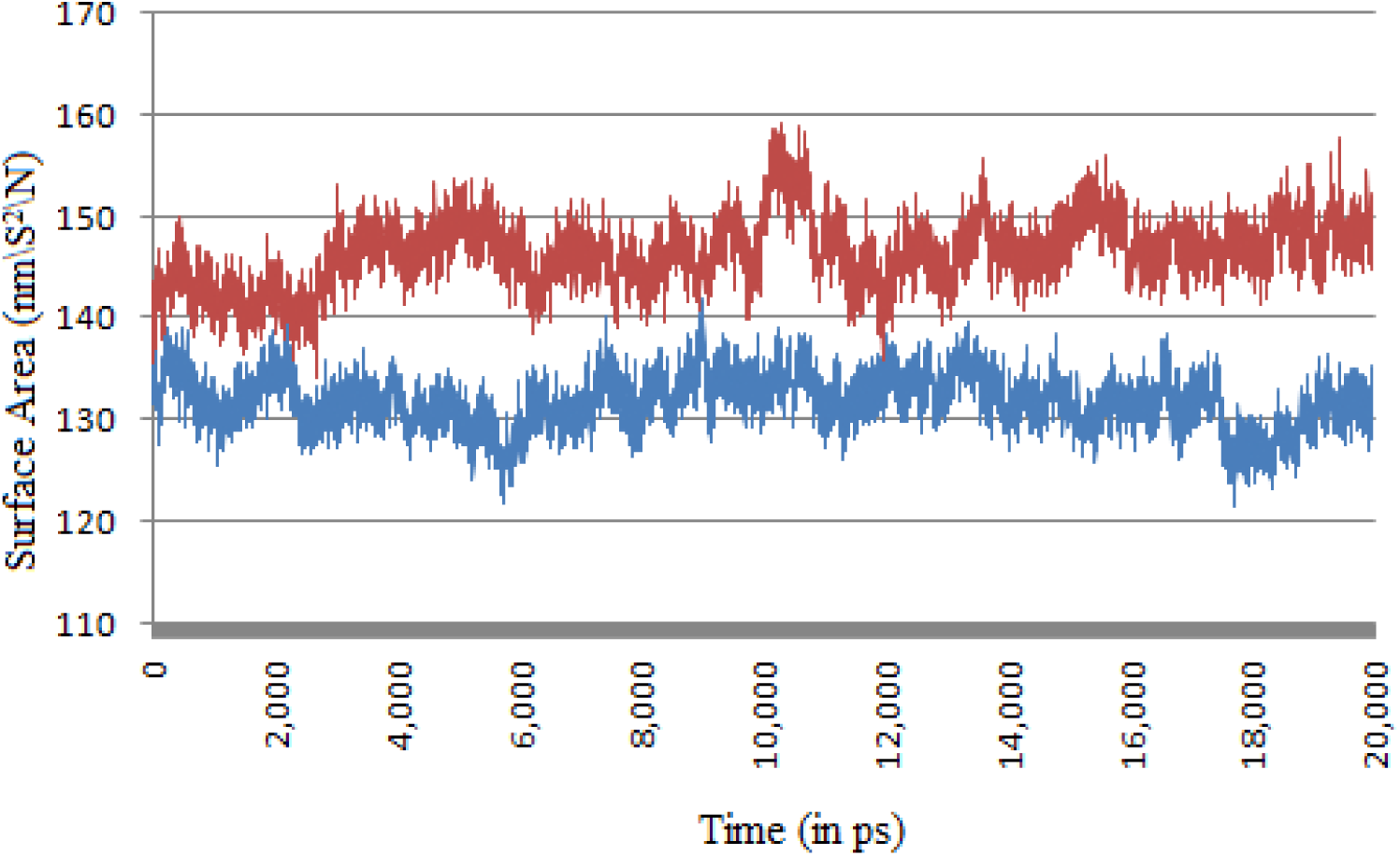
Total Solvant Accessible Surface Area of Focal Adhesion Kinase with (Blue) and without (Red) bound Solanesol.

**Figure 8:**
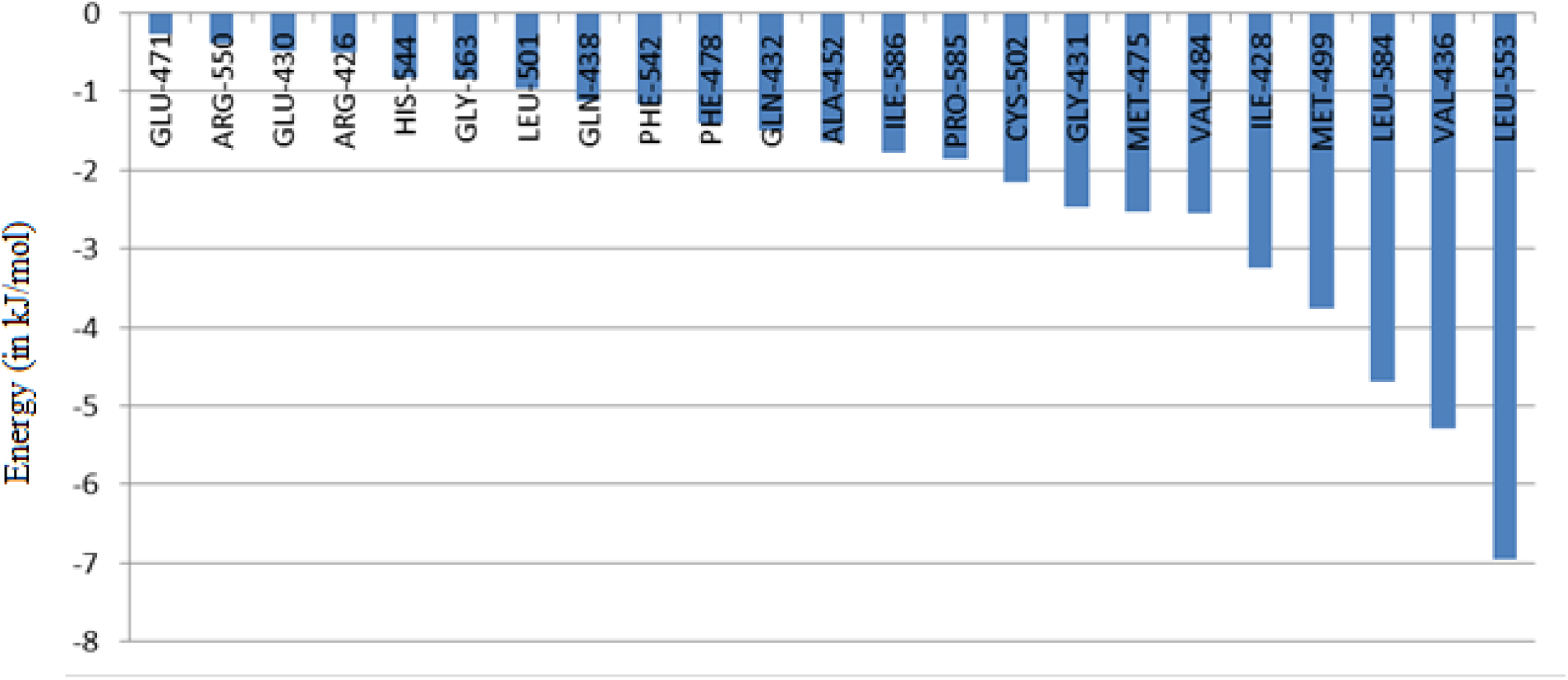
MM-PBSA based residue energy profile for active site residues.

**Figure 9:**
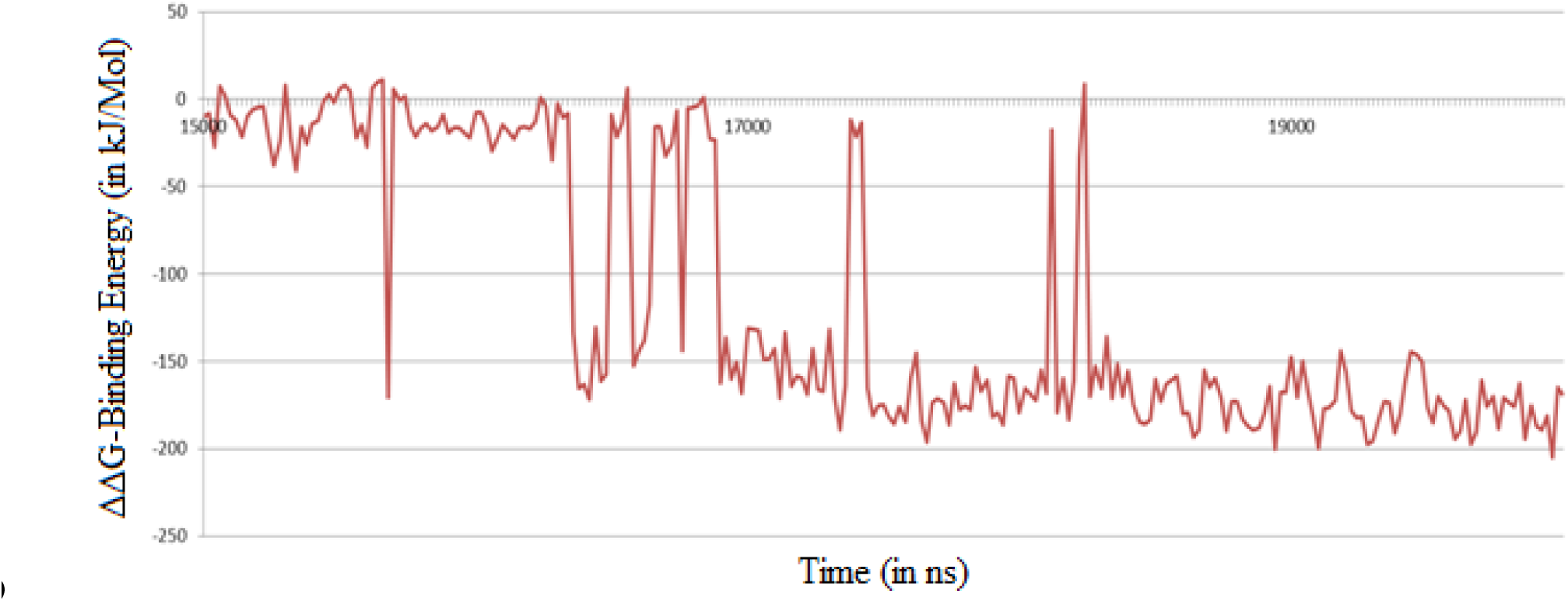
Binding Energy (ΔΔG) [kJ/mol]) using MM-PBSA method w.r.t time.

**Table 4:** MM-PBSA based final Binding free energy of Solanesol with Focal Adhesion Kinase

| $\Delta G$-vdw (in kJ/Mol) | $\Delta G$-electro (in kJ/Mol) | $\Delta G$-polar (in kJ/Mol) | $\Delta G$-SAS (in kJ/Mol) | $\Delta\Delta G$-BE (in kJ/Mol) |
| --- | --- | --- | --- | --- |
| -145.35 | -1.71 | 52.05 | -18.85 | **-113.85** |

## Conclusion

In this study, we have established a new method for scoring of binding pocket residues based on blind docking methodology of highly flexible residue. Knowledge based scoring function, MD simulation and binding free-energy calculations were performed to investigate and validate the binding conformation of Solanesol on Focal adhesion kinase at the molecular level found by LBCSR. This scoring was crucial as blind docking fails to give high confidence data for binding of highly flexible residues.

Based on the proposed LBCSR score, residues were identified which give high contact score with ligand in multiple blind docking instances. Refined docking was used to identify the best pose for this site. Knowledge based scoring function, DSX, was used to validate the docking result on which 20ns MD simulation was performed. Simulation of only protein structure of FAK protein was performed for establishing a control for MD simulations. The MM-PBSA binding free-energy calculations further confirmed the results of the MD simulation.

The MM-PBSA binding free-energy calculations further confirmed the results of the MD simulation. This analysis, established a correlation between the LBCSR and favorably interacting residues. It is in a good agreement with experimental results which shows Solanesol binds at ATP binding site and inhibit the phosphorylation of Focal Adhesion Kinase. This study may help in further analysis and understanding of flexible compounds and inhibition of FAK using the same.

## References

Cohen, L. A., & Guan, J. L. (2005). Mechanisms of focal adhesion kinase regulation. Current cancer drug targets, 5(8), 629–643.

Darden, T., York, D., & Pedersen, L. (1993). Particle mesh Ewald: An N · log (N) method for Ewald sums in large systems. The Journal of chemical physics, 98(12), 10089–10092.

Frisch, M., Trucks, G.W., Schlegel, H.B., Scuseria, G.E., Robb, M.A., Cheeseman, J.R., Scalmani, G., Barone, V., Mennucci, B., Petersson, G.A. and Nakatsuji, H., (2009). gaussian 09, Gaussian. Inc, Wallingford, CT.

George, D.M., Breinlinger, E.C., Friedman, M., Zhang, Y., Wang, J., Argiriadi, M., Bansal-Pakala, P., Barth, M., Duignan, D.B., Honore, P. and Lang, Q., (2014). Discovery of selective and orally bioavailable protein kinase Cθ (PKCθ) inhibitors from a fragment hit. Journal of medicinal chemistry, 58(1), 222–236.

Guan, J. L. (1997). Role of focal adhesion kinase in integrin signaling. The international journal of biochemistry & cell biology, 29(8), 1085–1096.

Hanahan, D., & Weinberg, R. A. (2011). Hallmarks of cancer: the next generation. cell,144(5), 646–674.

Hanahan, D., & Weinberg, R. A. (2000). The hallmarks of cancer cell 100: 57–70.

Hetényi, C., & van der Spoel, D. (2002). Efficient docking of peptides to proteins without prior knowledge of the binding site. Protein science, 11(7), 1729–1737.

Hetényi, C., & van der Spoel, D. (2006). Blind docking of drug-sized compounds to proteins with up to a thousand residues. FEBS letters, 580(5), 1447–1450.

Hess, B., Bekker, H., Berendsen, H. J., & Fraaije, J. G. (1997). LINCS: a linear constraint solver for molecular simulations. Journal of computational chemistry, 18(12), 1463–1472.

Hess, B., Kutzner, C., Van Der Spoel, D., & Lindahl, E. (2008). GROMACS 4: algorithms for highly efficient, load-balanced, and scalable molecular simulation. Journal of chemical theory and computation, 4(3), 435–447.

Kim, S., Thiessen, P.A., Bolton, E.E., Chen, J., Fu, G., Gindulyte, A., Han, L., He, J., He, S., Shoemaker, B.A. and Wang, J., 2015. PubChem substance and compound databases. Nucleic acids research, gkv951.

Kumari, R., Kumar, R., & Lynn, A. (2014). g_mmpbsa□ A GROMACS Tool for High-Throughput MM-PBSA Calculations. Journal of chemical information and modeling, 54(7), 1951–1962.

Lazebnik, Y. (2010). What are the hallmarks of cancer? Nature Reviews Cancer, 10(4), 232–233.

Massova, I., & Kollman, P. A. (2000). Combined molecular mechanical and continuum solvent approach (MM-PBSA/GBSA) to predict ligand binding. Perspectives in drug discovery and design, 18(1), 113–135.

Morris, G. M., Huey, R., Lindstrom, W., Sanner, M. F., Belew, R. K., Goodsell, D. S., & Olson, A. J. (2009). AutoDock4 and AutoDockTools4: Automated docking with selective receptor flexibility. Journal of computational chemistry, 30(16), 2785–2791.

National Center for Biotechnology Information (NCBI). PubChem Compound Database; CID=5477212, https://pubchem.ncbi.nlm.nih.gov/compound/5477212 (accessed Aug. 11, 2016).

Neudert, G., & Klebe, G. (2011). DSX: a knowledge-based scoring function for the assessment of protein–ligand complexes. Journal of chemical information and modeling, 51(10), 2731–2745.

Okamoto, Y., Tsuji, M., & Yamazaki, H. (1994). U.S. Patent No. 5,306,714. Washington, DC: U.S. Patent and Trademark Office.

Pettersen, E. F., Goddard, T. D., Huang, C. C., Couch, G. S., Greenblatt, D. M., Meng, E. C., & Ferrin, T. E. (2004). UCSF Chimera—a visualization system for exploratory research and analysis. Journal of computational chemistry, 25(13), 1605–1612.

Roe, S. J., Oldfield, M. F., Geach, N., & Baxter, A. (2013). A convergent stereocontrolled synthesis of [3-14C] solanesol. Journal of Labelled Compounds and Radiopharmaceuticals, 56(9-10), 485–491.

Severson, R. F., Ellington, J. J., Schlotzhauer, P. F., Arrendale, R. F., & Schepartz, A. I. (1977). Gas chromatographic method for the determination of free and total solanesol in tobacco. Journal of chromatography A, 139(2), 269–282.

Srivastava, S., Raj, K., Khare, P., Bhaduri, A.P., Chander, R., Raghubir, R., Mahendra, K., Rao, C.N. and Prabhu, S.K. (2009). Novel hybrid natural products derived from solanesol as wound healing agents. Indian journal of chemistry. Section B, Organic including medicinal, 48(2), 237.

Suzuki, H., Tomida, A., & Nishimura, T. (1990). Cytocidal Activity of a Synthetic Isoprenoid, N-Solanesyl-N, N′-bis (3, 4-dimethoxy-benzyl) ethylenediamine, and Its Potentiation of Antitumor Drugs against Multidrug-resistant and Sensitive Cells in vitro. Japanese journal of cancer research, 81(3), 298–303.

Tomida, A., & Suzuki, H. (1990). Synergistic Effect in Culture of Bleomycin-group Antibiotics and N-Solanesyl-N, N’-bis (3, 4-dimethoxybenzyl) ethylenediamine, a Synthetic Isoprenoid. Japanese journal of cancer research, 81(11), 1184–1190.

Van Aalten, D. M., Bywater, R., Findlay, J. B., Hendlich, M., Hooft, R. W., & Vriend, G. (1996). PRODRG, a program for generating molecular topologies and unique molecular descriptors from coordinates of small molecules. Journal of computer-aided molecular design, 10(3), 255–262.

Van Nimwegen, M. J., & van de Water, B. (2007). Focal adhesion kinase: a potential target in cancer therapy. Biochemical pharmacology, 73(5), 597–609.

Yan, N., Liu, Y., Gong, D., Du, Y., Zhang, H., & Zhang, Z. (2015). Solanesol: a review of its resources, derivatives, bioactivities, medicinal applications, and biosynthesis. Phytochemistry Reviews, 14(3), 403–417.

Zhao, C. J., Zu, Y. G., Li, C. Y., & Tian, C. Y. (2007). Distribution of solanesol in Nicotiana tabacum. Journal of forestry research, 18(1), 69–72.

